# Computational drug repositioning for the identification of new agents to sensitize drug-resistant breast tumors across treatments and receptor subtypes

**DOI:** 10.1101/2023.03.30.534178

**Authors:** Katharine Yu, Amrita Basu, Christina Yau, Denise M Wolf, Hani Goodarzi, Sourav Bandyopadhyay, James E Korkola, Gillian L Hirst, Smita Asare, I-SPY 2 TRIAL investigators, Angela Demichele, Nola Hylton, Douglas Yee, Laura Esserman, Laura van ’t Veer, Marina Sirota

## Abstract

Drug resistance is a major obstacle in cancer treatment and can involve a variety of different factors. Identifying effective therapies for drug resistant tumors is integral for improving patient outcomes. In this study, we applied a computational drug repositioning approach to identify potential agents to sensitize primary drug resistant breast cancers. We extracted drug resistance profiles from the I-SPY 2 TRIAL, a neoadjuvant trial for early stage breast cancer, by comparing gene expression profiles of responder and non-responder patients stratified into treatments within HR/HER2 receptor subtypes, yielding 17 treatment-subtype pairs. We then used a rank-based pattern-matching strategy to identify compounds in the Connectivity Map, a database of cell line derived drug perturbation profiles, that can reverse these signatures in a breast cancer cell line. We hypothesize that reversing these drug resistance signatures will sensitize tumors to treatment and prolong survival. We found that few individual genes are shared among the drug resistance profiles of different agents. At the pathway level, however, we found enrichment of immune pathways in the responders in 8 treatments within the HR+HER2+, HR+HER2-, and HR-HER2-receptor subtypes. We also found enrichment of estrogen response pathways in the non-responders in 10 treatments primarily within the hormone receptor positive subtypes. Although most of our drug predictions are unique to treatment arms and receptor subtypes, our drug repositioning pipeline identified the estrogen receptor antagonist fulvestrant as a compound that can potentially reverse resistance across 13/17 of the treatments and receptor subtypes including HR+ and triple negative. While fulvestrant showed limited efficacy when tested in a panel of 5 paclitaxel-resistant breast cancer cell lines, it did increase drug response in combination with paclitaxel in HCC-1937, a triple negative breast cancer cell line.

## 1 Introduction

Breast cancer is the most common cancer diagnosis in women worldwide and is expected to make up 15.3% of all new cancer cases in the United States in 2020^1^. While the prognosis for women with stage I or stage II breast cancer is excellent, 10-15% of newly diagnosed breast cancers are locally advanced cancers which have significantly poorer outcomes. Additionally, breast cancer is an incredibly heterogenous disease and research has shown that breast cancers with different molecular features can have different treatment responses^2,3^. Breast cancers can be stratified into receptor subtypes based on immunohistochemistry markers for ER, PR, and HER2, which are commonly used for therapeutic decision making^4^. Several of these receptor subtypes, which include triple negative, or ER-PR-HER2-tumors, and HER2+ tumors, represent patient populations with more aggressive disease even in early stage who could benefit from improved treatment^5^.

While breast cancer treatments have advanced, no treatment is effective in 100% of breast cancer patients. Drug resistance in cancer is a multi-faceted problem that involves a variety of biological determinants such as tumor heterogeneity, tumor burden and growth kinetics, physical barriers, the immune system, and the tumor microenvironment^6^. While there has been much research into understanding and overcoming drug resistance, it remains one of the largest challenges in cancer today and new approaches are needed to tackle this problem.

The I-SPY 2 TRIAL (Investigation of Serial studies to Predict Your Therapeutic Response with Imaging And molecular anaLysis 2) is an adaptive phase II clinical trial of neoadjuvant treatment for women with high risk, locally advanced breast cancer^7 8 9 10 11 12^. The trial uses an adaptive design to accelerate the clinical trial process with the goal of identifying optimal treatment regimens for patient subsets based on HR, HER2, and MammaPrint^5^. While the I-SPY 2 trial has been successful in graduating numerous drugs, patients who fail to respond to the neoadjuvant treatments in the trial tend to have worse outcomes^13 14^. Identifying more efficacious treatments for these non-responder patients with primary drug resistance may improve patient outcomes.

We applied a computational drug repurposing approach to identify potential agents to include in the trial for patients unlikely to respond to agent classes tested in the trial to date. Drug repurposing offers advantages over traditional drug development by greatly reducing development costs and providing shorter paths to approval, as drug safety has already been established during the drug’s original regulatory process. Our group has previously developed and applied a computational drug repositioning approach which involves generating a disease gene expression signature by comparing disease samples to control samples, and then identifying a drug that can reverse this disease signature^17^. Potential drug hits can be found by using datasets such as the Connectivity Map (CMap) and the Library of Integrated Network-Based Cellular Signatures (L1000) which have generated thousands of drug perturbation expression profiles. This gene expression based computational drug repurposing approach has previously been used to identify effective treatments for a number of different indications, including several cancer types^18 19^. It has also been used to predict agents to reverse drug resistance in acute lymphoblastic leukemia and non-small cell lung cancer ^20 21^.

In this study, we leveraged the I-SPY2-990 mRNA/RPPA data compendium^22^ to extract drug resistance signatures by comparing the pre-treatment expression profiles of responders to non-responders within each receptor subtype and treatment arm. We then applied a computational drug repositioning approach to identify agents which can reverse these primary drug resistance signatures, and experimentally tested the top drug hit in a panel of paclitaxel-resistant breast cancer cell lines. This is the first large scale attempt to apply this transcriptomics-based drug repositioning pipeline to the receptor subtypes of breast cancer.

## 2 Methods

### 2.1 I-SPY2 Gene Expression and Clinical Data

I-SPY 2 is a multicenter, phase II adaptive clinical trial for women with high-risk stage II/III breast cancer. Patients are classified into receptor subtypes based on hormone-receptor (HR), HER2, and MammaPrint status and assigned to one of several investigational therapies or the control regimen using an adaptive randomization engine which gives greater weight to treatments with a higher estimated response rate in the patient’s tumor subtype. The primary endpoint is pathologic complete response (pCR, no residual invasive disease in breast or nodes) at the time of surgery. The analysis is modified intention to treat and patients who do not proceed to surgery, withdraw from the trial, or receive non-protocol therapy are considered non-pCR.

We used pre-treatment biopsy samples from the closed arms of the ISPY2 trial (n=990), which were assayed using custom Agilent array designs (15746 and 32627). Normalized data for each array was generated by centering the log2 transformed gMeanSignal of all probes within the array to the 75^th^ percentile of all probes. A fixed value of 9.5 was added to avoid negative values. Genes with multiple probes were averaged and ComBat was applied to adjust for platform-biases^22^.

We define drug resistant patients as patients with Residual Cancer Burden (RCB) III measured at time of surgery and drug sensitive patients as patients with RCB 0 or I at time of surgery. While we initially included RCB II patients in the drug resistant group, we removed the RCB II patients in our final analysis to achieve better separation in predictive signals distinguishing responders and non-responders. We kept receptor subtype and treatments with at least three patients in the resistant and sensitive groups, resulting in 19 receptor subtype-treatment pairs.

### 2.2 Differential Expression to Identify Drug Resistance Genes

We used limma to perform differential expression between the drug resistant and drug sensitive samples within treatments and receptor subtypes. We then filtered the differential expression results by p-value and log-fold change to generate the resistance gene lists. We chose a p-value threshold of 0.01 because the differences between the resistant and sensitive tumors were relatively subtle and very few genes met the typical q-value cutoff of 0.05. To identify the optimal log fold change cutoff for each differential expression gene list, we selected the log fold change value that best separated the drug resistant and drug sensitive samples after filtering for p-value < 0.01. Specifically, we iterated over a range of potential log2 fold change cutoffs (start = 1, end = 0, step size = 0.1) and applied k-means clustering (k=2) at each cutoff to identify two clusters of samples. We then calculated the Mathew’s correlation coefficient (MCC) to evaluate how well the k-means derived clusters match the actual clinical labels of drug resistant and drug sensitive samples. We used the log2 fold change cutoff with the highest MCC value to generate our drug resistance gene lists. Only drug resistance gene lists with a sufficient number of genes (>50) were kept for further analysis.

### 2.3 Gene Set Enrichment Analysis

For the GSEA analysis, the drug resistance profiles were ranked by their log fold-change values. We used the fgsea R package^37^ to calculate normalized enrichment scores (NES) and FDR values from these ranked lists. The NES reflects the degree to which a gene set is overrepresented at the top or bottom of the ranked list of genes (the enrichment score) divided by the mean enrichment score for all dataset permutations. Normalizing the enrichment score allows for comparison across gene sets. We downloaded the 50 Hallmark gene sets from the MSigDB Collections^38^.

### 2.4 Computational Drug Repositioning

We applied our previously published drug repositioning pipeline^17^ to identify potential therapeutics to reverse drug resistance in breast cancer patients. At a high level, the method works by identifying drugs that have reversed differential gene expression profiles compared to the drug resistance profile. We hypothesize that reversing the expression patterns of drug resistance genes will drive the tumor towards a drug sensitive state.

To prioritize drugs that have the potential to reverse the drug resistance genes, we used drug perturbation profiles from CMap V2, which includes 6100 profiles consisting of 1309 distinct chemical compounds. We applied a filtering step previously described by Chen et al. (2017) to keep high quality drug perturbation profiles. We further subset this dataset to include only drug profiles that were generated using MCF-7, the only breast cancer cell line in CMap, resulting in a final dataset of 756 profiles.

Our drug repositioning pipeline uses a non-parametric, rank-based pattern-matching strategy based on the Kolmogorov-Smirnov (KS) statistic to assess the enrichment of drug resistance genes in a ranked drug perturbation gene list. We calculate a reverse gene expression score (RGES) of each drug by matching resistance gene expression and drug gene expression using the KS test. Significance of the score is assessed by comparing with scores generated from 100,000 random permutations, and further corrected by the multiple hypothesis test. FDR < 0.05 was used to select drug hits.

### 2.5 Validation experiments for fulvestrant

To validate fulvestrant as a compound to overcome drug resistance, we first selected paclitaxel-resistant breast cancer cell lines because paclitaxel was used as the standard therapy in the ISPY2 trial. We selected three paclitaxel-resistant and three paclitaxel-sensitive cell lines from Daemen et al. (2015) from within the HR+HER2- and HR-HER2-receptor subtypes. Daemen et al. only identified 2 Paclitaxel-sensitive cell lines and 2 Paclitaxel-resistant cell lines for the HR+HER2+ subtype, so we included all four HR+HER2+ cell lines in our validation experiment. Additionally, since Daemen et al. did not identify any Paclitaxel-resistant HR-HER2+ cell lines in their study, we did not include any HR-HER2+ cell lines in our validation experiment.

We ordered 16 cell lines from ATCC (Table 3) which were recovered using the cell media recommended for each cell line by ATCC. We failed to culture three cell lines: MDA-MB-134-VI, BT-483, UACC-812. Cell line density was determined by seeding cell lines at the following densities (625, 1250, 2500, 5000, 10000, 20000) and then monitoring their growth curves for 72 hours. For the drug treatment experiments, the cell lines were seeded at the optimal density determined in the previous cell line density experiments and incubated overnight before treatment. For the single agent experiments, the cell lines were treated in triplicate with a top dose of 10uM in 1:3 dilutions for a total of 12 doses with paclitaxel (Sigma-Aldrich Product Number T7191), fulvestrant (Sigma-Aldrich Product Number I4409), and staurosporine which was used as a positive control. After 72hr, cell line viability was measured using the CellTiter-Glo Luminescent Cell Viability Assay following the manufacturer’s instructions. For the sequential treatment experiments, 1uM of fulvestrant was added to each well 6 hours before treatment with paclitaxel. The 1 uM dose and 6 hour time point were chosen based on the dose and time point used to generate the CMAP profile for fulvestrant. For the combination treatment experiments, the cell lines were treated with paclitaxel as described above in combination with 10uM fulvestrant.

## 3 Results

### 3.1 Study design and datasets

In this study, we applied our drug repositioning pipeline to the drug resistance signatures derived from the I-SPY2 trial (Figure 1). Pre-treatment samples from ∼990 patients in 9 experimental arms of the trial and concurrent controls were profiled using the Agilent 44K array, as previously described^22^. The clinical data for these samples includes the HR/HER2 receptor subtype of each sample, treatment, and treatment response including pathologic complete response (pCR), defined as the absence of invasive cancer in the breast and lymph nodes, and residual cancer burden (RCB) information. RCB scores are a continuous variable based on the primary tumor dimensions, the cellularity in the tumor bed, and the axillary nodal burden after neoadjuvant therapy. The continuous RCB score can then be divided into discrete RCB classes (0, 1, 2, 3) based on predefined cutoffs^23^. An RCB of 0 indicates pathologic complete response while an RCB of 1-3 indicates increasing amounts of residual cancer. 109 samples were missing RCB information and excluded from the analysis. The data used in this study form part of the ISPY2-990 mRNA/RPPA data compendium^22^ recently deposited on GEO (GSE196096). A summary of the clinical data, including receptor subtype which we define by the HR and HER2 status of the tumor, is provided in Supplementary Table 1 and the corresponding arm for each treatment is provided in Supplementary Table 2.

**Figure 1.**
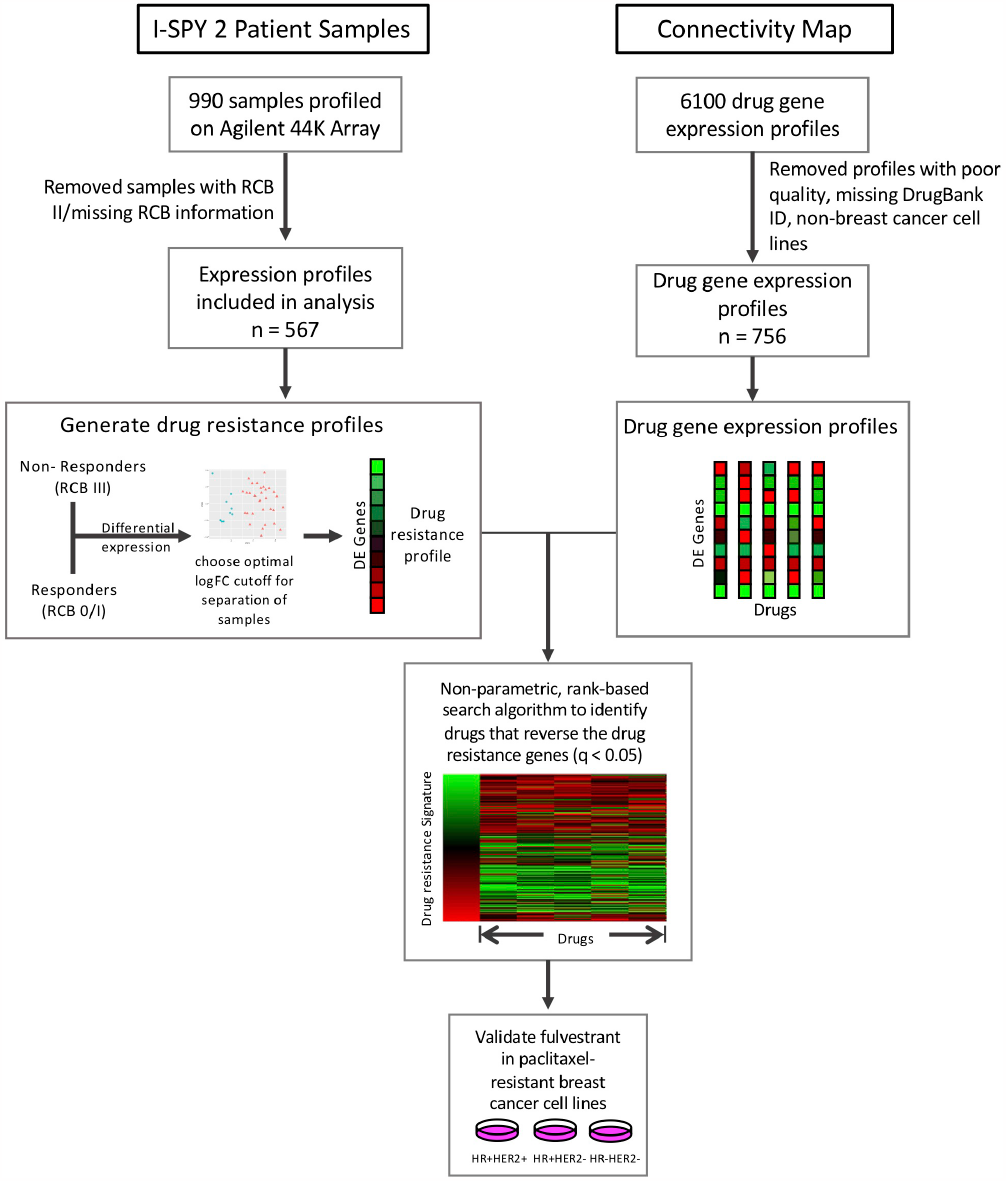
Study overview. Drug resistance gene lists were generated for each subtype and treatment arm by performing differential expression between responders (RCB 0/I) and non-responders (RCB III). We then compared these drug resistance gene profiles to the Connectivity Map drug perturbation profiles for the MCF7 breast cancer cell line to identify drugs that can reverse these drug resistance genes. We tested our top hit, fulvestrant, in paclitaxel-resistant breast cancer cell lines.

### 3.2 Drug resistance gene profiles overlap at the pathway level and include previously implicated drug resistance genes

We first classified each pre-treatment biopsy sample from the ISPY 2 trial as drug sensitive or drug resistant using the RCB class from the clinical data. We define drug sensitive tumors as having an RCB of 0 or I and we define drug resistant tumors as having an RCB of III. While we originally defined resistant tumors as having RCB II or III, we found a more distinct signal when resistance is defined using RCB III only and RCB II tumors are removed from the data set (Supplementary Table 3 and Supplementary Figure 1).

We performed differential expression analysis between drug sensitive and drug resistant patients within individual treatments, by receptor subtype. We analyzed only the receptor subtype-treatment pairs with a minimum of 3 samples in both the drug sensitive group and the drug resistant group, which resulted in a total of 19 subtype-treatment pairs (Table 1). Of note, there was an insufficient number of HR-HER2+ tumors for our within-treatment analysis and this receptor subtype was excluded from our study.

**Table 1.** Summary of receptor subtype and treatments.

| Treatment | Receptor subtype | Sensitive | Resistant | # of genes in resistance profile |
| --- | --- | --- | --- | --- |
| Paclitaxel + ABT 888 + Carboplatin | HR+HER2- | 28 | 4 | 109 |
| Paclitaxel + ABT 888 + Carboplatin | HR+ HER2- | 10 | 7 | 182 |
| Paclitaxel + AMG 386 | HR- HER2- | 30 | 5 | 55 |
| Paclitaxel + AMG 386 | HR+ HER2- | 19 | 13 | 165 |
| Paclitaxel + Ganetespib | HR- HER2- | 24 | 4 | 124 |
| Paclitaxel + Ganetespib | HR+ HER2- | 12 | 9 | 85 |
| Paclitaxel | HR- HER2- | 31 | 9 | 69 |
| Paclitaxel | HR+ HER2- | 22 | 23 | 531 |
| Paclitaxel + MK-2206 | HR- HER2- | 18 | 3 | 201 |
| Paclitaxel + MK-2206 | HR+ HER2- | 7 | 7 | 593 |
| Paclitaxel + Neratinib | HR- HER2- | 16 | 6 | 146 |
| Paclitaxel + Neratinib | HR+ HER2- | 3 | 3 | 147 |
| Paclitaxel + Neratinib | HR+ HER2+ | 17 | 7 | 88 |
| Paclitaxel + Pembrolizumab | HR+ HER2- | 17 | 7 | 217 |
| Paclitaxel + Pertuzumab + Trastuzumab | HR+ HER2+ | 12 | 3 | 170 |
| Paclitaxel + Trastuzumab | HR+ HER2+ | 7 | 3 | 176 |
| T-DM1 + Pertuzumab | HR+ HER2+ | 19 | 4 | 157 |

**Table 2.** Table of genes in drug resistance profiles.

| **Gene Symbol** | **# of drug resistance profiles** | **Description** | **References** |
| --- | --- | --- | --- |
| POU2AF1 | 5 | Transcriptional coactivator |  |
| SERPINA3 | 5 | Member of the serpin family of proteins | 24 39 |
| EPHX2 | 4 | Member of the epoxide hydrolase family | 40 |
| STC2 | 4 | Secreted, homodimeric glycoprotein | 25 |
| CHST8 | 4 | Member of the sulfotransferase 2 family |  |
| CXCL11 | 4 | CXC chemokine, chemotactic for interleukin-activated T-cells | 41 |
| HAPLN3 | 4 | Member of the hyaluronan and proteoglycan binding link protein gene family | 42 |
| CXCL13 | 4 | CXC chemokine, lymphocyte B chemoattractant | 43 |
| EVL | 4 | Actin-associated proteins | 44 |
| HSD11B1 | 4 | Microsomal enzyme, reversibly catalyzes conversion of cortisol to cortisone |  |
| IDO1 | 4 | Heme enzyme, catalyzes tryptophan catabolism | 45 |
| IL21R | 4 | Cytokine receptor for interleukin 21 | 46 |
| SEL1L3 | 4 | Protein coding gene |  |
| SLC22A5 | 4 | Organic cation and sodium-dependent high affinity carnitine transporter | 47 |
| TNFRSF17 | 4 | Receptor for TNFSF13B/BLyS/BAFF and TNFSF13/APRIL |  |
| ZBED2 | 4 | Transcriptional regulator | 48 |
| ANKRD22 | 4 | Protein coding gene |  |
| LPPR3 | 4 | Member of the lipid phosphate phosphatase (LPP) family |  |

We generated drug resistance gene profiles for each receptor subtype and treatment by filtering the differential expression analysis results by p-value (0.01) and then selecting the optimal log-fold change cutoff to achieve maximal separation between the drug resistant and drug sensitive tumors (see Methods). Drug resistance gene profiles with fewer than 50 genes were removed as we had previously found this to be the minimum sufficient number of genes required for the drug repositioning pipeline^18^. The drug resistance gene profiles for the remaining 17 receptor subtype-treatment pairs are included in Supplementary Data 1. We also generated a more general drug resistance profile by comparing all resistant tumors to all sensitive samples while adjusting for receptor subtype and treatments, but this profile achieved poor separation of resistant and sensitive tumors (Supplementary Figure 2).

We found that few individual genes are shared across the receptor subtype and treatment drug resistance gene profiles (Figure 2A). However, of the 18 genes that appear in at least 4 of the subtype-treatment pair resistance profiles, 11 have been implicated in drug resistance or drug response based on the literature. For example, SERPINA3, which was present in five of the drug resistance gene profiles, including paclitaxel with neratinib and paclitaxel with pembrolizumab in the HR+HER2-subtype, has been implicated in drug resistance in TNBC cells^24^. Additionally, STC2, which has been implicated in drug resistance in cervical cancer^25^, was in the following four drug resistance gene profiles: paclitaxel in the HR+HER2-subtype, paclitaxel with ganetespib in the HR+HER2-subtype, paclitaxel with pertuzumab and trastuzumab in the HR+HER2+ subtype, and paclitaxel with trastuzumab in the HR+HER2+ subtype.

**Figure 2.**
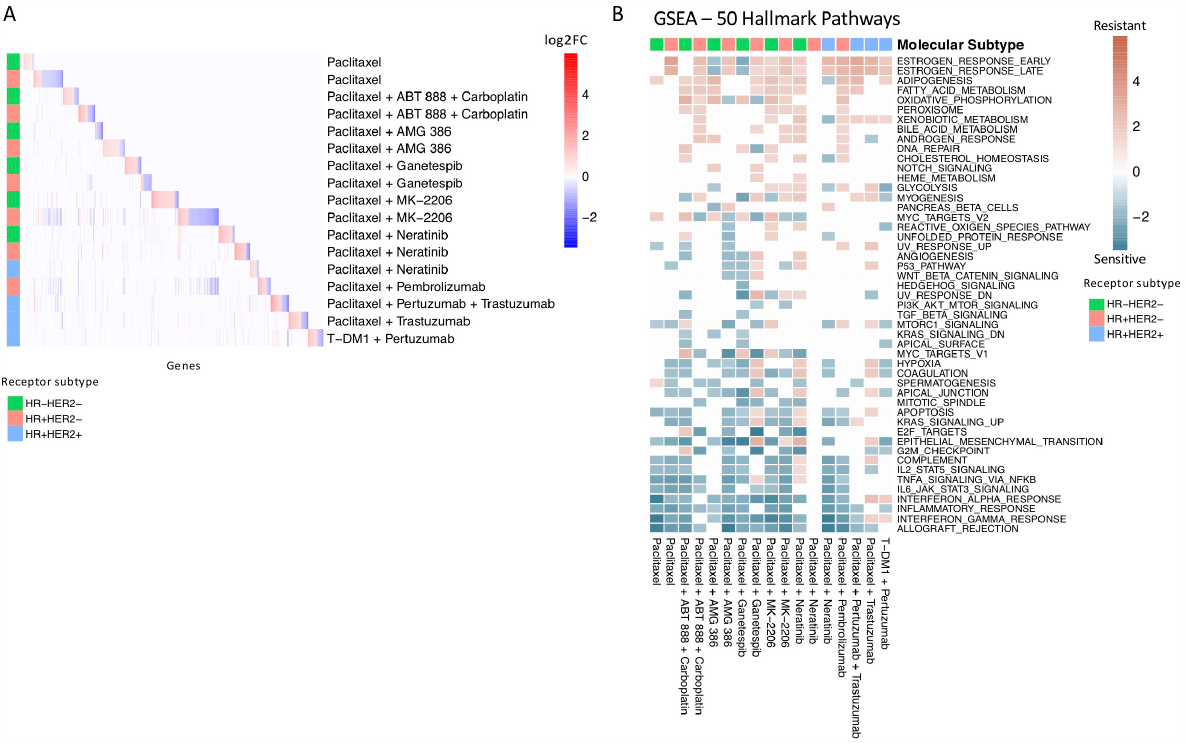
Drug resistance gene profiles overlap at pathway level. **(A)** Heatmap of significant differentially expressed genes in each treatment and receptor subtype arm. The colored annotation bar on the left side of the heatmap indicates the receptor subtype of the treatment arm. The colors within the heatmap indicates log-fold change with red indicating significantly upregulated genes and blue indicating significantly downregulated genes. White indicates that a gene was not differentially expressed in the specific treatment and receptor subtype arm. **(B)** Gene Set Enrichment Analysis of drug resistance signatures in treatment and molecular subtype arms using MsigDB’s 50 hallmark pathways. Red boxes indicate enrichment in non-responders and turquoise boxes indicate enrichment in responders. Significant (q-value < 0.05) normalized enrichment scores (NES) are shown.

We then performed Gene Set Enrichment Analysis (GSEA)^26^ to investigate the differences between the drug sensitive and drug resistant tumors at the pathway level with the 50 hallmark pathways from MSigDB (Figure 2B). Similar to previous studies^27 28^, we found an enrichment of immune pathways in drug sensitive tumors compared to drug resistant tumors in 14 out of the 17 receptor subtype and treatment pairs, including as expected the HR+HER2-subtype in the pembrolizumab treatment. We also found an enrichment of estrogen response pathways in drug resistant tumors in 12 of the receptor subtype-treatment pairs, 10 of which are in the hormone-receptor positive receptor subtypes. The estrogen response pathway has also been previously implicated in chemoresistance^29^.

### 3.3 Prediction of drug sensitizing agents based on expression identifies fulvestrant as a potential therapeutic

We applied a transcriptomics-based drug repositioning pipeline^17^ to compare the drug resistance gene profiles to the Connectivity Map, a public dataset of drug perturbation profiles, in order to identify compounds which have the reversed differential gene expression profiles compared to the drug resistance gene profiles. We hypothesize that if we can identify a drug which can downregulate the genes that are upregulated in drug resistance and upregulate the genes which are downregulated in drug resistance, then this drug may induce chemosensitivity in resistant breast cancer tumors. Out of 756 high quality gene perturbation profiles in the Connectivity Map dataset derived from a breast cancer cell line, the median number of significant drug hits (q-value < 0.05 and RES < 0) per receptor subtype-treatment pair was 49 (min: 1, max: 256). The drug hits for each receptor subtype and treatment are reported in Supplementary Data 2.

Although the number of individual genes that overlap across the drug resistance gene profiles of the different receptor subtype-treatment pairs was limited, we observed 22 drugs that appeared as hits in at least 9/17 of the drug resistance gene profiles (Figure 3A and Supplementary Figure 3).

**Figure 3.**
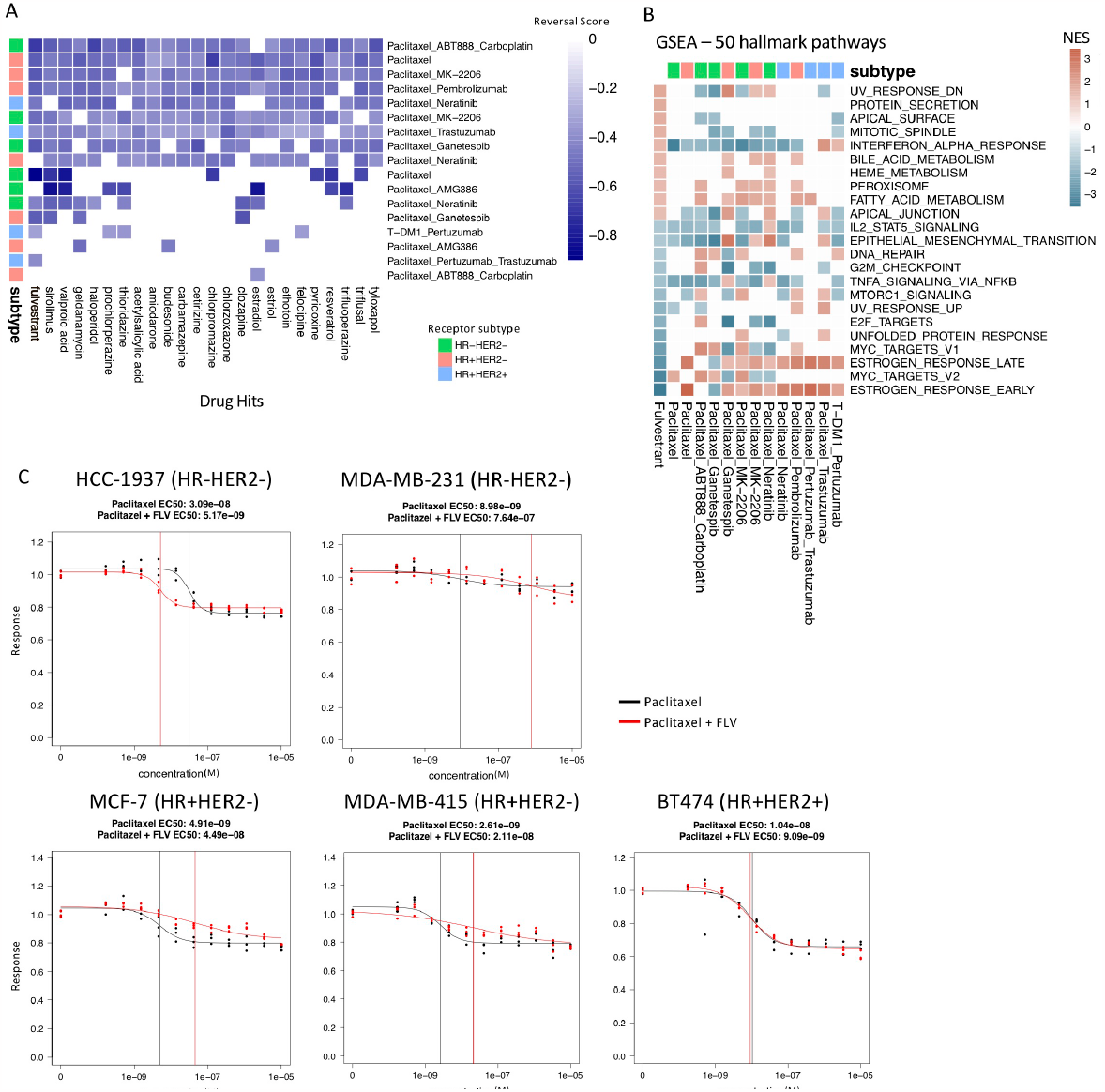
Drug hits and validation experiments. **(A)** Heatmap of the 22 most common drug hits (q-value < 0.05 and RES < 0) across treatment and molecular subtype arms. Color indicates strength of reversal score and white color indicates that drug is not a significant hit in the specific treatment and molecular subtype arm. **(B)** GSEA analysis comparing fulvestrant perturbation profile (first column) to the drug resistance profiles using MsigDB’s 50 hallmark pathways. Only pathways that have significant NES scores (q-value < 0.05) in the fulvestrant perturbation profile are shown. **(C)** Drug response of paclitaxel alone (black) and fulvestrant and paclitaxel in combination (red) tested in paclitaxel-resistant breast cancer cell lines. The vertical lines indicate the EC50 values. Fulvestrant and paclitaxel given in combination increases response in the HCC-1937 cell line.

Of note, we identified fulvestrant as a drug hit that significantly reversed 13/17 of the drug resistance profiles. It is predicted to reverse the drug resistance profiles in 5/6 treatment groups for TN; 4/4 for HR+HER2+; and 4/7 for HR+HER2- (Figure 3A). Fulvestrant is a selective estrogen receptor degrader used in the treatment of hormone-receptor positive and HER2-advanced breast cancer in post-menopausal woman who have not previously been treated with endocrine therapy. We performed GSEA on the fulvestrant drug perturbation signature from the Connectivity Map to investigate the pathways which are reversed by fulvestrant and examined the enrichment of these pathways in the drug resistance profiles (Figure 3B). Unsurprisingly, fulvestrant seems to downregulate the estrogen response pathways and cell cycle pathways. A previous study also showed that fulvestrant may reverse drug resistance in multidrug-resistant breast cancer cell lines independent of estrogen receptor expression^30^. For these reasons, we selected fulvestrant for further validation experiments.

### 3.4 Fulvestrant validation experiments demonstrate limited efficacy in breast cancer cell lines

In order to validate fulvestrant as a drug candidate that can reverse drug resistance, we first needed to identify a panel of drug-resistant breast cancer cell lines. We selected cell lines that are resistant to paclitaxel because paclitaxel is a standard therapy in the I-SPY 2 trial. The Daemen et al. study screened 90 experimental and approved drugs, including paclitaxel, in a panel of 70 breast cancer cell lines. Based on the drug response data from this study, we selected paclitaxel-resistant and paclitaxel-sensitive breast cancer cell lines within each receptor subtype. The cell lines selected for the validation experiments are listed in Table 3 and were ordered from ATCC. We were unable to grow three of the cell lines (MDA-MB-134-VI, BT-483, UACC-812), which were excluded from the drug response experiments.

**Table 3.** Summary of breast cancer cell line responses to paclitaxel.

| Cell line | Receptor Subtype | **−log<sub>10</sub>(EC<sub>50</sub>)** | Paclitaxel status |
| --- | --- | --- | --- |
| HCC-1937 | HR-HER2- | 5.24 | Resistant* |
| MDA-MB-231 | HR-HER2- | 5.46 | Resistant |
| MCF-7 | HR+HER2- | 6.77 | Resistant* |
| MDA-MB-415 | HR+HER2- | 6.83 | Resistant |
| BT-474 | HR+HER2+ | 7.44 | Resistant |
| MDA-MB-436 | HR-HER2- | 7.69 | Sensitive |
| BT-549 | HR-HER2- | 7.99 | Sensitive* |
| HCC-38 | HR-HER2- | 8.11 | Sensitive* |
| MDA-MB-361 | HR+HER2+ | 8.15 | Sensitive |
| ZR-751 | HR+HER2- | 8.26 | Sensitive |
| HCC-1143 | HR-HER2- | 8.56 | Sensitive |
| T-47D | HR+HER2- | 8.84 | Sensitive* |
| ZR-7530 | HR+HER2+ | 9.48 | Sensitive |

\*Indicates that Paclitaxel response matches response in the Daemen et al. paper

Next, we treated the breast cancer cell lines with paclitaxel to validate the drug responses from the Daemen et al. study^31^. We used the mean EC50 response as the cutoff to separate the resistant and sensitive cell lines. We identified five cell lines that were resistant to paclitaxel based on this cutoff, two of which were also found to be resistant in the Daemen et al. study (Table 3). The discrepancy between our drug responses and the drug responses in the Daemen et al. study may be due in part to the different drug response metrics that were used. The Daemen et al. study used GI50 while we used EC50 to measure drug response. Out of the five cell lines that we determined to be resistant to paclitaxel, two were HR-HER2-, two were HR+HER2-, and one was HR+HER2+.

We then tried two different treatment strategies for testing fulvestrant in the paclitaxel resistant cell lines. In the first treatment strategy, we treated the paclitaxel resistant cell lines with fulvestrant for 6 hours before adding paclitaxel. This sequential treatment approach gives the cell lines time to become sensitized by fulvestrant before being treated with paclitaxel. This sequential treatment approach (Supplementary Figure 4) did not result in a change in response to paclitaxel in the paclitaxel-resistant cell lines. In the second treatment strategy, we treated the paclitaxel-resistant cell lines with both fulvestrant and paclitaxel in combination for 72 hours.

Out of the five paclitaxel-resistant cell lines, this combination treatment strategy resulted in an increase in response in one cell line, HCC-1937, with an EC50 shift from 3.09e-8 to 5.17e-9 M, and a decrease in sensitivity in MCF-7 and MDA-MB-415 (Figure 3C). Interestingly, HCC-1937 is a triple negative breast cancer cell line, suggesting perhaps an estrogen receptor independent mechanism of action.

## 4 Discussion

Drug resistance is the primary factor that limits cures in cancer patients. In this study, we applied a computational drug repositioning approach to identify potential FDA-approved agents for patients with primary drug-resistant tumors in the I-SPY 2 trial.

We generated drug resistance profiles for each receptor subtype and treatment by comparing the expression profiles of responder to non-responder patients. While we were unable to identify genes that were present across every drug resistance profile, many of the genes which appeared in multiple drug resistance profiles have been previously implicated in drug resistance. SERPINA3, which was upregulated in multiple drug resistance profiles, has been shown to reduce sensitivity of TNBC cells to cisplatin upon overexpression^24^. Similarly, STC2, which was also upregulated in multiple drug resistance profiles, has been found to be significantly elevated in cisplatin resistant cervical cancer cells^25^. We were able find literature support for a number of genes that were present in multiple drug resistance profiles, suggesting that our drug resistance profiles are capturing aspects of known biology about drug resistance.

When we performed gene set enrichment analysis on the drug resistance profiles, we identified enrichment of estrogen response and metabolic pathways in resistant tumors compared to sensitive tumors. This is in line with previous studies which have shown that estrogen can promote resistance to chemotherapeutic drugs in ER+ human breast cancer cells through regulation of the Bcl-2 proto-oncogene^29^. Unsurprisingly, the estrogen response pathways were primarily enriched in the HR+ groups in our analysis. Previous studies have also shown that metabolic pathways are key mediators of drug resistance in breast cancer. Fatty acid metabolism, which was enriched in resistant tumors across multiple receptor subtype and treatments in our analysis, has previously been implicated in drug resistance through mechanisms such as increased fatty acid oxidation, which can generate energy for cancer cells, or decreased membrane fluidity, which can affect drug uptake^32^. Oxidative phosphorylation was also found to be enriched across multiple receptor subtype and treatments, similar to previous studies which have shown that tamoxifen-resistant MCF-7 breast cancer cells display increased levels of oxidative phosphorylation^33^.

We identified potential drug candidates by searching for drugs in the CMAP dataset that can significantly reverse these drug-resistance profiles. Fulvestrant was our most common drug hit and it was predicted to significantly reverse 85% of the drug resistance profiles. An *in vitro* study using multi-drug resistant breast cancer cell lines showed that fulvestrant can induce sensitivity to doxorubicin^30^. Interestingly, they found that this response was independent of the ER status of the breast cancer cell lines and may involve an interaction with P-glycoprotein. Sirolimus, also known as rapamycin, was another drug that appeared across multiple drug resistance profiles. Previous studies have shown that sirolimus may enhance the effects of chemotherapies in breast cancer cell lines^34^ and osteosarcoma cell lines^35^. Additionally, MK-2206 targets the same pathway and was shown to be effective in the I-SPY 2 trial^9^. While we selected fulvestrant to test in vitro because it appeared as a hit in the greatest number of drug resistance profiles, the other drug hits may be promising candidates for reversing drug sensitivity in breast cancer.

For the validation experiments, we first selected breast cancer cell line that were either sensitive or resistant to paclitaxel based on the Daemen et al. study (2015). We then validated the drug responses by treating these cell lines with paclitaxel and we identified five cell lines that are paclitaxel-resistant. We treated these paclitaxel-resistant breast cancer cell lines with fulvestrant and paclitaxel, both sequentially and in combination. While fulvestrant showed limited efficacy in a majority of the cell lines, fulvestrant in combination with paclitaxel did increase drug response in one triple negative cell line, HCC-1937, suggesting the potential of fulvestrant as a combination treatment for drug-resistant tumors within specific genetic contexts. It is worth noting, however, that the HR+HER2-cell lines did not respond to fulvestrant, which was unexpected, especially since one of the cell lines, MCF-7, was used to generate the CMap drug perturbation profiles used for prediction. It is possible that a higher dose or a longer pre-treatment time with fulvestrant may be necessary to induce a response in these cell lines. Alternatively, these cell lines may reflect hormone receptor-positive tumors that do not respond to chemotherapy, as identified in previous clinical trials^36^.

Our study has several limitations which we discuss here. First, the drug perturbation data used to make the predictions was derived from MCF-7, a single HR+HER2-cell line. Had the drug perturbation data included multiple breast cancer cell lines spanning the different receptor subtypes, the predictions may have been improved. Second, the primary tumor expression profiles from the I-SPY 2 study are from pre-treatment samples only. Thus, the drug resistance profiles that we generated primarily reflect intrinsic drug resistance rather than adaptive drug resistance, the latter of which would require post-treatment samples. Additionally, after stratifying the I-SPY 2 patient samples by receptor subtype and treatment, the number of samples within some groups were relatively small, limiting the power of the study. Similarly, our validation experiments were performed in a limited number of breast cancer cell lines. Future experiments should incorporate more patient samples, including post-treatment samples, to generate more robust drug resistance profiles to inform predictions, which should be based on more diverse cell lines that better capture breast cancer heterogeneity. We also hope to test additional drug hits in a larger panel of breast cancer cell lines, such as the panel used in Daemen et. al, to better understand the genomic context contributing to drug response.

In summary, we used a computational drug repurposing approach to identify potential agents to sensitize drug resistant breast cancers. We generated drug resistance profiles for each receptor subtype and treatment in the I-SPY 2 trial and found that estrogen response and metabolic pathways are enriched in resistant tumors and immune pathways are enriched in sensitive tumors. We then compared these drug resistance profiles to the drugs in CMAP and identified drug hits for each resistance profile. We tested fulvestrant in a panel of five paclitaxel-resistant breast cancer cell lines and found that it increased drug response in combination with paclitaxel in the cell line HCC-1937.

## Acknowledgements

With support from Quantum Leap Healthcare Collaborative (I-SPY 2 trial sponsor), NCI grant PO1-CA210961, Breast Cancer Research Foundation (BCRF-20-165), NIH/NCI CCMI (Grant U54CA274502-01), NIH/NCI CCSG (Grant P30-CA82103), Breast Cancer Research – Atwater Trust

## Author Contributions

KY, MS, LV, AB, CY and DW designed the study. JEK, GH, LS, I-SPY 2 TRIAL investigators, AD, NH, DY, LE were involved in data generation. KY performed the computational analysis. All authors contributed to interpretation of the results. LV, HG and SB assisted with planning the validation experiments. KY and MS wrote the manuscript. KY, MS, AB, CY, DW, GH, DY, LV edited the paper. MS and LV supervised the work. All authors reviewed and approved the manuscript.

## Declaration of Interests

CY reports institutional research funding from the National Institutes of Health (NIH); reports financial support from Quantum Leap Healthcare Collaborative; reports patents: U.S Provisional Application No. 63/314,065, U.S Provisional Application No. 63/341,579 and US Application No. 18/174,191; and served on the Data Safety Monitoring Board for the ISPY2 Trial. DW reports institutional research funding from the National Cancer Institute and Quantum Leap Healthcare Collaborative. SB reports institutional research funding and consulting fees from Revolution Medicines; and is an employee of and holds stock in Rezo Therapeutics. JK is a co-founder and holds stock in Convergent Genomics. GH reports research grants: P01CA210961 and U01CA196406; and spouse holds stock in Moderna, Exact Sciences, Gilead and Nanostring. NH reports institutional research funding and grants from the National Institutes of Health (NIH). DY reports institutional research funding from Quantum Leap Healthcare Collaborative and the National Institutes of Health / National Cancer Institute (P01 CA210961-01A1); reports consulting fees from Martell Diagnostics and reports honoraria and travel support from PER for speaking at the International Breast Cancer Conference. LE reports institutional research funding from Merck; reports participation as a Medical Advisory Panel Member for Blue Cross Blue Shield; and is a website author of UpToDate. LVV is co-founder of Agendia NV. MS reports consulting fees from Exxagen; and holds stock in Aria Pharmaceuticals and Somnics. All other authors declare no competing interests.

## Data Availability

All data used in this study are publicly available. ISPY2-990 mRNA/RPPA data compendium is available on GEO (GSE196096). The ConnectivityMap data is available at https://clue.io/.

## References

1. National Institutes of Health; National Cancer Institute. Surveillance, Epidemiology, and End Results Program. Cancer stat facts: female breast cancer. https://seer.cancer.gov/statfacts/html/breast.html.

2. Blows, F. M. et al. Subtyping of Breast Cancer by Immunohistochemistry to Investigate a Relationship between Subtype and Short and Long Term Survival: A Collaborative Analysis of Data for 10,159 Cases from 12 Studies. PLoS Med 7, e1000279 (2010).

3. Valero, V. V., Buzdar, A. U. & Hortobagyi, G. N. Locally Advanced Breast Cancer. Oncologist 1, 8–17 (1996).

4. Vallejos, C. S. et al. Breast cancer classification according to immunohistochemistry markers: subtypes and association with clinicopathologic variables in a peruvian hospital database. Clin Breast Cancer 10, 294–300 (2010).

5. Barker, A. et al. I-SPY 2: An Adaptive Breast Cancer Trial Design in the Setting of Neoadjuvant Chemotherapy. Clinical Pharmacology & Therapeutics 86, 97–100 (2009).

6. Vasan, N., Baselga, J. & Hyman, D. M. A view on drug resistance in cancer. Nature 575, 299–309 (2019).

7. Rugo, H. S. et al. Adaptive Randomization of Veliparib–Carboplatin Treatment in Breast Cancer. N Engl J Med 375, 23–34 (2016).

8. Park, J. W. et al. Adaptive Randomization of Neratinib in Early Breast Cancer. N Engl J Med 375, 11–22 (2016).

9. Chien, A. J. et al. MK-2206 and Standard Neoadjuvant Chemotherapy Improves Response in Patients With Human Epidermal Growth Factor Receptor 2-Positive and/or Hormone Receptor-Negative Breast Cancers in the I-SPY 2 Trial. J Clin Oncol 38, 1059–1069 (2020).

10. Nanda, R. et al. Effect of Pembrolizumab Plus Neoadjuvant Chemotherapy on Pathologic Complete Response in Women With Early-Stage Breast Cancer: An Analysis of the Ongoing Phase 2 Adaptively Randomized I-SPY2 Trial. JAMA Oncol 6, 676–684 (2020).

11. Clark, A. S. et al. Neoadjuvant T-DM1/pertuzumab and paclitaxel/trastuzumab/pertuzumab for HER2+ breast cancer in the adaptively randomized I-SPY2 trial. Nat Commun 12, 6428 (2021).

12. Pusztai, L. et al. Durvalumab with olaparib and paclitaxel for high-risk HER2-negative stage II/III breast cancer: Results from the adaptively randomized I-SPY2 trial. Cancer Cell 39, 989–998.e5 (2021).

13. Yee, D. et al. Association of Event-Free and Distant Recurrence–Free Survival With Individual-Level Pathologic Complete Response in Neoadjuvant Treatment of Stages 2 and 3 Breast Cancer. JAMA Oncol 6, 1–9 (2020).

14. Spring, L. M. et al. Pathological complete response after neoadjuvant chemotherapy and impact on breast cancer recurrence and survival: a comprehensive meta-analysis. Clin Cancer Res 26, 2838–2848 (2020).

15. von Minckwitz, G. et al. Trastuzumab Emtansine for Residual Invasive HER2-Positive Breast Cancer. New England Journal of Medicine 380, 617–628 (2019).

16. Masuda, N. et al. Adjuvant Capecitabine for Breast Cancer after Preoperative Chemotherapy. New England Journal of Medicine 376, 2147–2159 (2017).

17. Sirota, M. et al. Discovery and preclinical validation of drug indications using compendia of public gene expression data. Sci Transl Med 3, 96ra77 (2011).

18. Chen, B. et al. Reversal of cancer gene expression correlates with drug efficacy and reveals therapeutic targets. Nat Commun 8, 16022 (2017).

19. Chen, B. et al. Computational Discovery of Niclosamide Ethanolamine, a Repurposed Drug Candidate That Reduces Growth of Hepatocellular Carcinoma Cells In Vitro and in Mice by Inhibiting Cell Division Cycle 37 Signaling. Gastroenterology 152, 2022–2036 (2017).

20. Spijkers-Hagelstein, J. a. P., Pinhanços, S. S., Schneider, P., Pieters, R. & Stam, R. W. Chemical genomic screening identifies LY294002 as a modulator of glucocorticoid resistance in MLL-rearranged infant ALL. Leukemia 28, 761–769 (2014).

21. Yeh, C.-T. et al. Trifluoperazine, an antipsychotic agent, inhibits cancer stem cell growth and overcomes drug resistance of lung cancer. Am J Respir Crit Care Med 186, 1180–1188 (2012).

22. Wolf, D. M. et al. Redefining breast cancer subtypes to guide treatment prioritization and maximize response: Predictive biomarkers across 10 cancer therapies. Cancer Cell 40, 609–623.e6 (2022).

23. Symmans, W. F. et al. Measurement of residual breast cancer burden to predict survival after neoadjuvant chemotherapy. J Clin Oncol 25, 4414–4422 (2007).

24. Zhang, Y. et al. Overexpression of SERPINA3 promotes tumor invasion and migration, epithelial-mesenchymal-transition in triple-negative breast cancer cells. Breast Cancer 28, 859–873 (2021).

25. Wang, Y., Gao, Y., Cheng, H., Yang, G. & Tan, W. Stanniocalcin 2 promotes cell proliferation and cisplatin resistance in cervical cancer. Biochem Biophys Res Commun 466, 362–368 (2015).

26. Subramanian, A. et al. Gene set enrichment analysis: A knowledge-based approach for interpreting genome-wide expression profiles. Proceedings of the National Academy of Sciences 102, 15545–15550 (2005).

27. Zitvogel, L., Galluzzi, L., Smyth, M. J. & Kroemer, G. Mechanism of action of conventional and targeted anticancer therapies: reinstating immunosurveillance. Immunity 39, 74–88 (2013).

28. Bracci, L., Schiavoni, G., Sistigu, A. & Belardelli, F. Immune-based mechanisms of cytotoxic chemotherapy: implications for the design of novel and rationale-based combined treatments against cancer. Cell Death Differ 21, 15–25 (2014).

29. Jiang, Z. et al. The role of estrogen receptor alpha in mediating chemoresistance in breast cancer cells. J Exp Clin Cancer Res 31, 42 (2012).

30. Huang, Y., Jiang, D., Sui, M., Wang, X. & Fan, W. Fulvestrant reverses doxorubicin resistance in multidrug-resistant breast cell lines independent of estrogen receptor expression. Oncol Rep 37, 705–712 (2017).

31. Daemen, A. et al. Modeling precision treatment of breast cancer. Genome Biology 14, R110 (2013).

32. Chen, M. & Huang, J. The expanded role of fatty acid metabolism in cancer: new aspects and targets. Precis Clin Med 2, 183–191 (2019).

33. Fiorillo, M., Sotgia, F., Sisci, D., Cappello, A. R. & Lisanti, M. P. Mitochondrial “power” drives tamoxifen resistance: NQO1 and GCLC are new therapeutic targets in breast cancer. Oncotarget 8, 20309–20327 (2017).

34. Zhang, J. et al. Rapamycin Antagonizes BCRP-Mediated Drug Resistance Through the PI3K/Akt/mTOR Signaling Pathway in mPRα-Positive Breast Cancer. Frontiers in Oncology 11, (2021).

35. Zhou, Y. et al. Sirolimus induces apoptosis and reverses multidrug resistance in human osteosarcoma cells in vitro via increasing microRNA-34b expression. Acta Pharmacol Sin 37, 519–529 (2016).

36. Albain, K. S. et al. Adjuvant chemotherapy and timing of tamoxifen in postmenopausal patients with endocrine-responsive, node-positive breast cancer: a phase 3, open-label, randomised controlled trial. The Lancet 374, 2055–2063 (2009).

37. Korotkevich, G. et al. Fast gene set enrichment analysis. bioRxiv 060012 (2021) doi:10.1101/060012.

38. Liberzon, A. et al. The Molecular Signatures Database (MSigDB) hallmark gene set collection. Cell Syst 1, 417–425 (2015).

39. Jinawath, N. et al. Oncoproteomic Analysis Reveals Co-Upregulation of RELA and STAT5 in Carboplatin Resistant Ovarian Carcinoma. PLOS ONE 5, e11198 (2010).

40. Vainio, P. et al. Arachidonic Acid Pathway Members PLA2G7, HPGD, EPHX2, and CYP4F8 Identified as Putative Novel Therapeutic Targets in Prostate Cancer. Am J Pathol 178, 525–536 (2011).

41. Zhang, Y. et al. CXCL11 promotes self-renewal and tumorigenicity of α2δ1+ liver tumor-initiating cells through CXCR3/ERK1/2 signaling. Cancer Lett 449, 163–171 (2019).

42. Heterogeneous Mechanisms of Secondary Resistance and Clonal Selection in Sarcoma during Treatment with Nutlin | PLOS ONE. https://journals.plos.org/plosone/article?id=10.1371/journal.pone.0137794.

43. Zhang, G. et al. CXCL-13 Regulates Resistance to 5-Fluorouracil in Colorectal Cancer. Cancer Res Treat 52, 622–633 (2020).

44. Padilla-Rodriguez, M. et al. The actin cytoskeletal architecture of estrogen receptor positive breast cancer cells suppresses invasion. Nat Commun 9, 2980 (2018).

45. Okamoto, A. et al. Indoleamine 2,3-dioxygenase serves as a marker of poor prognosis in gene expression profiles of serous ovarian cancer cells. Clin Cancer Res 11, 6030–6039 (2005).

46. Mittal, D. et al. Improved Treatment of Breast Cancer with Anti-HER2 Therapy Requires Interleukin-21 Signaling in CD8+ T Cells. Cancer Res 76, 264–274 (2016).

47. Okabe, M. et al. Profiling SLCO and SLC22 genes in the NCI-60 cancer cell lines to identify drug uptake transporters. Mol Cancer Ther 7, 3081–3091 (2008).

48. Hyter, S. et al. Developing a genetic signature to predict drug response in ovarian cancer. Oncotarget 9, 14828–14848 (2017).

