## Supplementary Materials for "Computational drug repositioning for the identification of new agents to sensitize drug-resistant breast tumors across treatments and receptor subtypes"

Supplementary Figure 1. Removing RCB II samples improves separation of drug sensitive and resistant samples

A

Drug resistance profile genes generated with RCB II samples

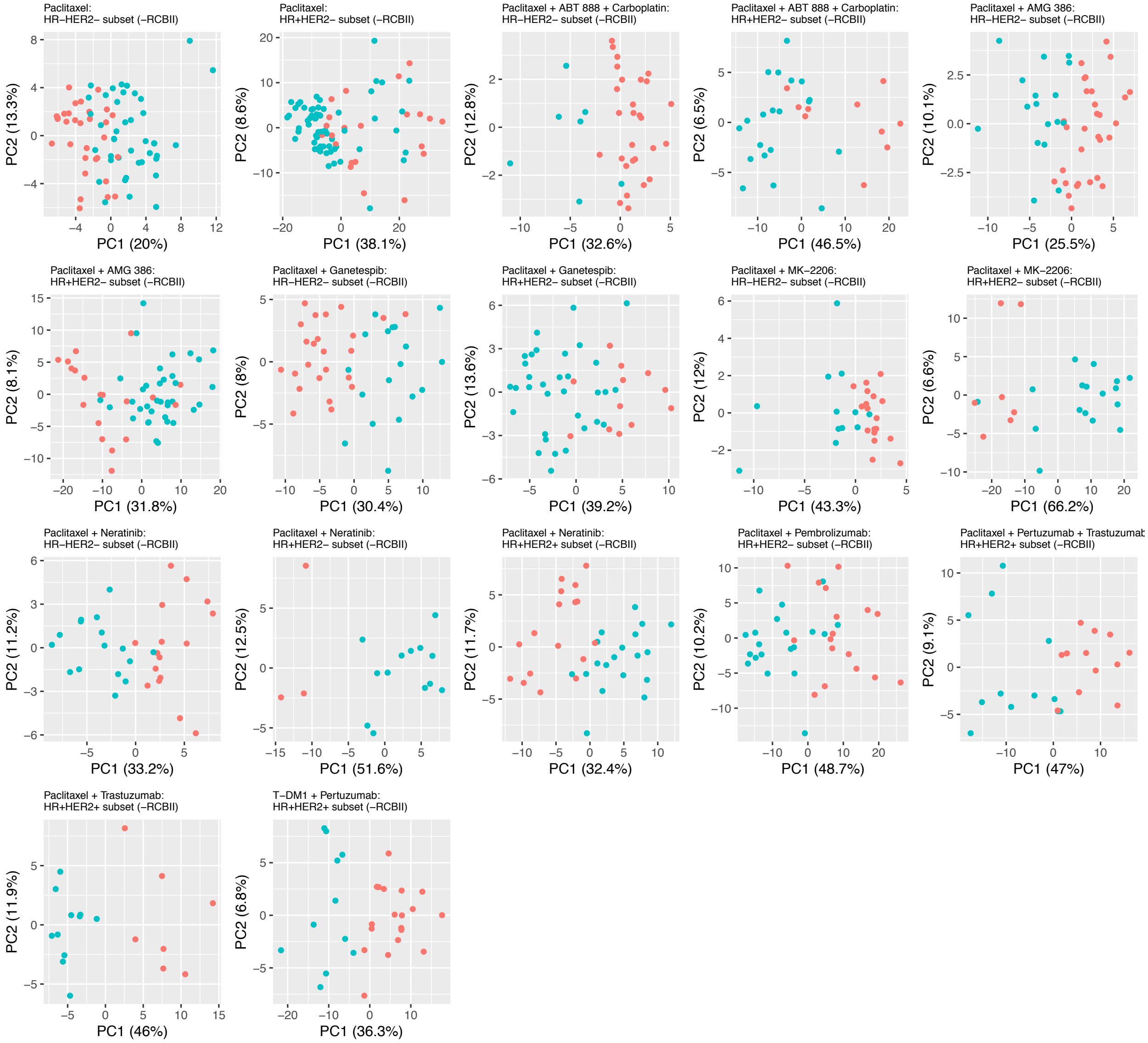

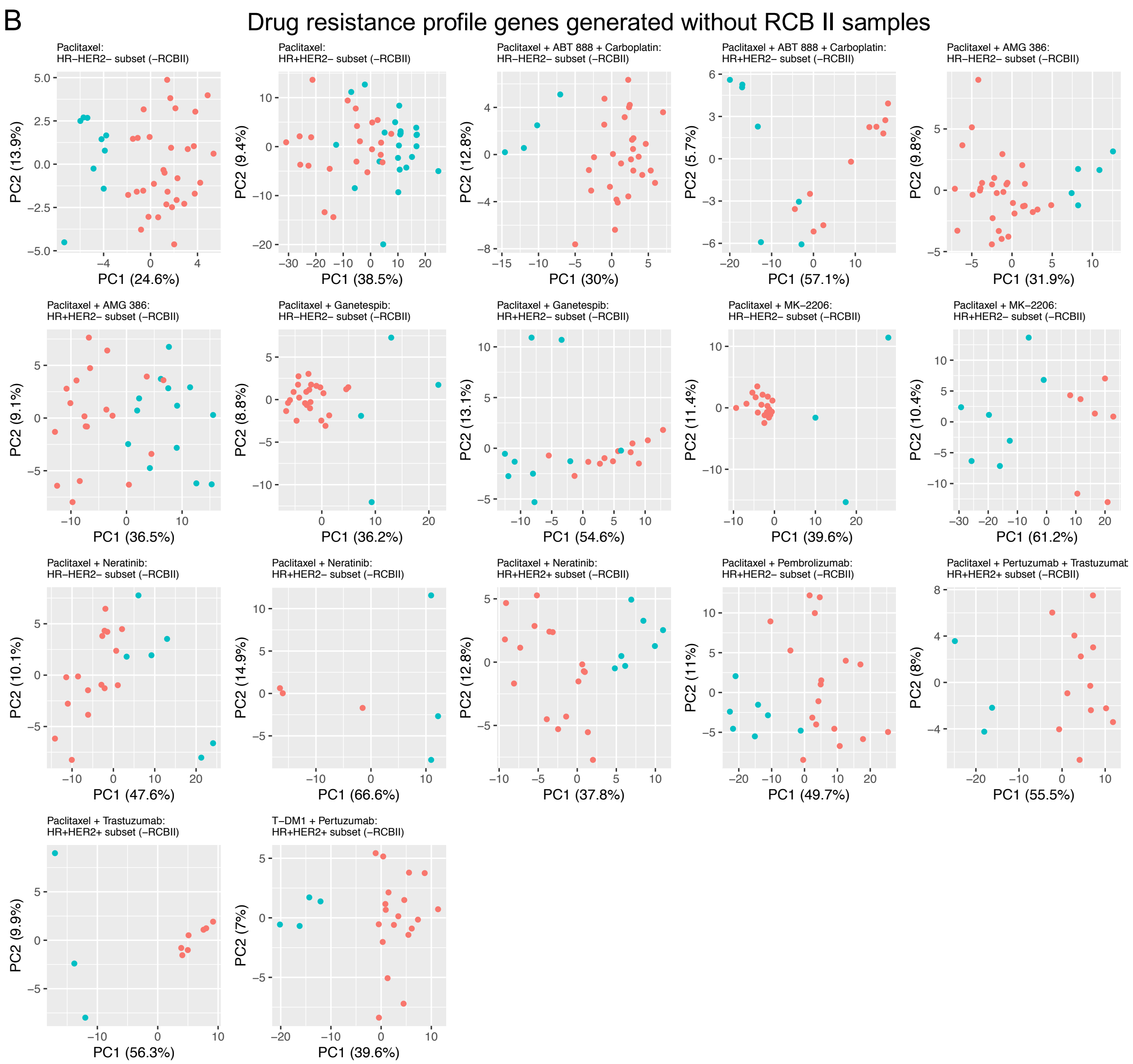


A. PCA plots of drug resistant and sensitive samples using drug resistance profile genes generated with RCB II samples. The x-axis shows principal component 1 with variance explained in parenthesis and y-axis shows principal component 2 with variance explained in parenthesis. Drug resistant samples are indicated by the red points and drug sensitive samples are indicated by the turquoise points. B. Same as the PCA plots in subfigure A but with RCB II samples removed. Removing the RCB II samples improves the separation of the drug resistant and drug sensitive samples in 13/17 of the molecular subtype and treatment arm pairs.

Supplementary Figure 2. Drug resistance profile using all samples

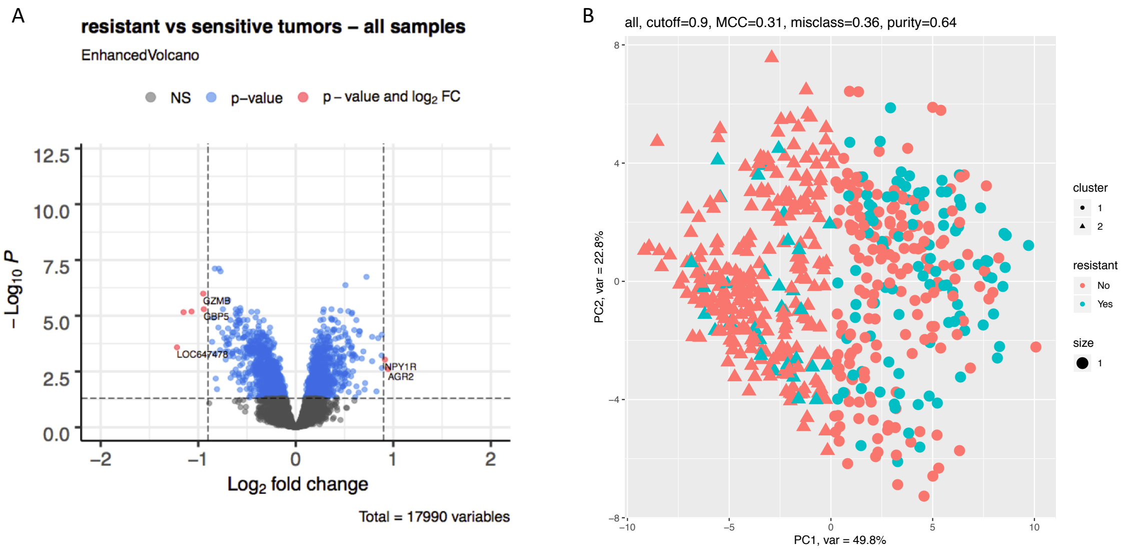


A. Volcano plot of differential expression results comparing all resistant tumors to all sensitive tumors (adjusted for molecular subtype and treatment arm). B. PCA plot showing the optimal log-fold change cutoff for the differential expression analysis results using all sensitive and resistant samples. Matthew's Correlation Coefficient (MCC) is 0.31, suggesting poor separation of resistant and sensitive samples.

Supplementary Figure 3. All drug hits across molecular subtype and treatment arms

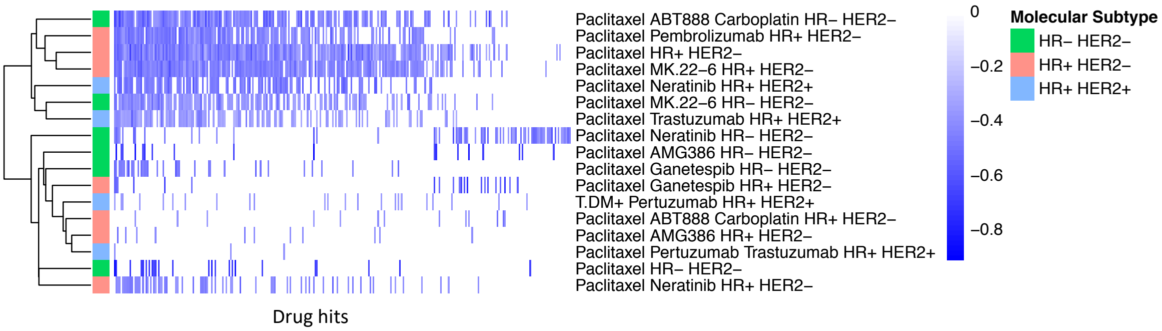


Heatmap of all drug hits (q-value < 0.05 and RES < 0) across treatment and molecular subtype arms. Color indicates strength of reversal score and white color indicates that drug is not a significant hit in specific treatment and molecular subtype arm.

Supplementary Figure 4. Cell line responses to sequential fulvestrant and paclitaxel treatment

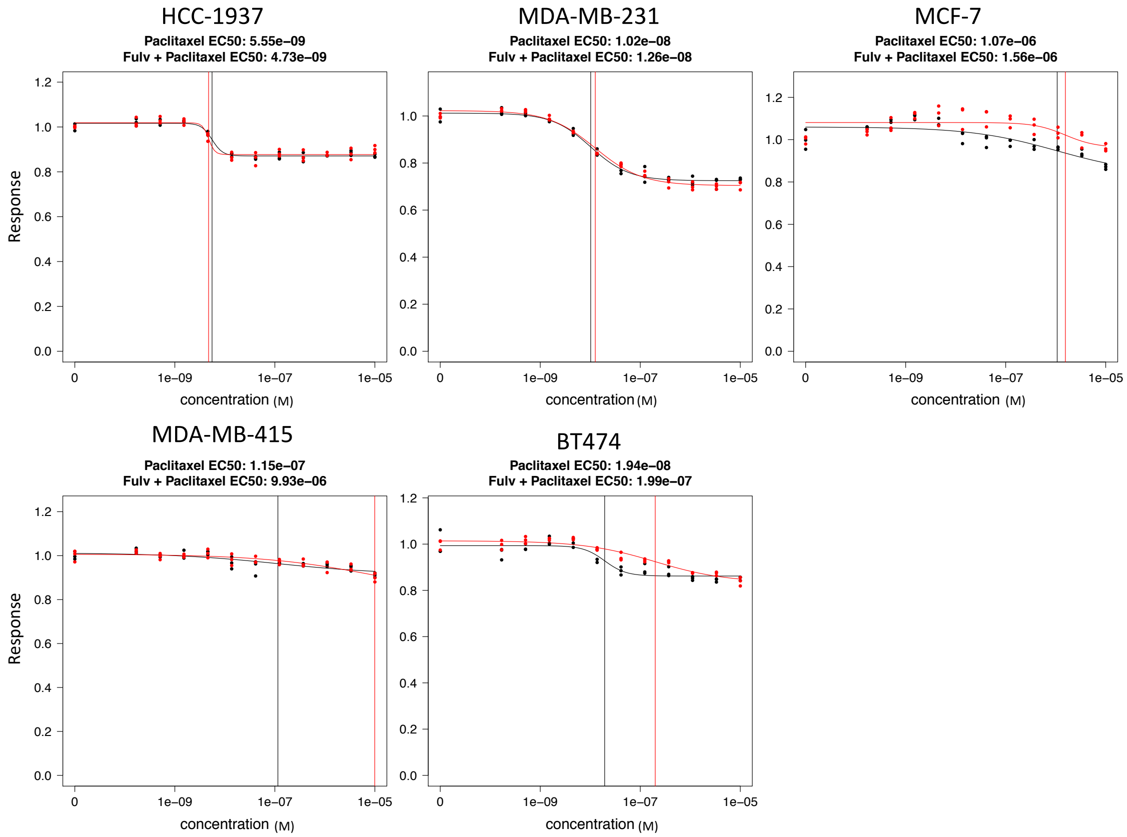


Paclitaxel-resistant breast cancer cell lines were treated with fulvestrant for 6 hours before treating with paclitaxel for 72 hours. Paclitaxel alone (black) and fulvestrant pre-treatment with paclitaxel (red) EC50's are indicated by the vertical colored lines. Fulvestrant pre-treatment does not seem to affect cell line response to paclitaxel.

Supplementary Table 1. Summary of I-SPY2 clinical data

| Treatment | HR+HER2- | HR+HER2+ | HR-HER2+ | HR-HER2- | Treatment total |
|---|---|---|---|---|---|
| Paclitaxel + ABT 888 + Carboplatin | 33 | 0 | 0 | 39 | **72** |
| Paclitaxel + Neratinib | 18 | 42 | 23 | 32 | **115** |
| Paclitaxel + Trastuzumab | 0 | 19 | 12 | 0 | **31** |
| Paclitaxel + AMG 386 | 62 | 0 | 0 | 53 | **115** |
| Paclitaxel | 95 | 0 | 0 | 84 | **179** |
| Paclitaxel + MK-2206 + Trastuzumab | 0 | 16 | 18 | 0 | **34** |
| Paclitaxel + Ganetespib | 48 | 0 | 0 | 45 | **93** |
| Paclitaxel + Pembrolizumab | 38 | 0 | 0 | 29 | **67** |
| T-DM1 + Pertuzumab | 0 | 35 | 17 | 0 | **52** |
| Paclitaxel + Ganitumab | 58 | 0 | 0 | 48 | **106** |
| Paclitaxel + MK-2206 | 28 | 0 | 0 | 32 | **60** |
| Paclitaxel + Pertuzumab + Trastuzumab | 0 | 29 | 15 | 0 | **44** |
| Paclitaxel + AMG 386 + Trastuzumab | 0 | 15 | 4 | 0 | **19** |
| **Total** | **380** | **156** | **89** | **362** | **987** |

| RCB | HR+HER2- | HR+HER2+ | HR-HER2+ | HR-HER2- | RCB Total |
|---|---|---|---|---|---|
| 0 | 64 | 49 | 52 | 141 | **306** |
| I | 44 | 21 | 8 | 54 | **127** |
| II | 160 | 44 | 11 | 95 | **310** |
| III | 76 | 20 | 4 | 35 | **135** |
| NULL | 36 | 22 | 14 | 37 | **109** |
| **Total** | **380** | **156** | **89** | **362** | **987** |

Supplementary Table 2. Mapping of treatments to arms in the ISPY-2 TRIAL.

| **Treatment** | **Arm** |
|---|---|
| Paclitaxel + ABT 888 + Carboplatin | veliparib/carboplatin |
| Paclitaxel + Neratinib | neratinib |
| Paclitaxel + Trastuzumab | control |
| Paclitaxel + AMG 386 | trebananib |
| Paclitaxel | control |
| Paclitaxel + MK-2206 + Trastuzumab | MK2206 |
| Paclitaxel + Ganetespib | genetespib |
| Paclitaxel + Pembrolizumab | pembrolizumab |
| T-DM1 + Pertuzumab | TDM1/pertuzumab |
| Paclitaxel + Ganitumab | ganitumab |
| Paclitaxel + MK-2206 | MK2206 |
| Paclitaxel + Pertuzumab + Trastuzumab | pertuzumab |
| Paclitaxel + AMG 386 + Trastuzumab | trebananib |

Supplementary Table 3. Removing RCB II increases the Mathew's Correlation Coefficient (MCC) of most molecular subtype and treatment arms

| **Treatment** | **Molecular Subtype** | **MCC with RCB II** | **MCC without RCB II** |
|---|---|---|---|
| Paclitaxel | HR-HER2- | 0.68 | 1 |
| Paclitaxel | HR+HER2- | 0.33 | 0.52 |
| Paclitaxel + ABT888 + Carboplatin | HR-HER2- | 0.91 | 1 |
| Paclitaxel + ABT888 + Carboplatin | HR+HER2- | 0.61 | 1 |
| Paclitaxel + AMG386 | HR+HER2- | 0.72 | 0.75 |
| Paclitaxel + AMG386 | HR-HER2- | 0.75 | 0.9 |
| Paclitaxel + Ganetespib | HR+HER2- | 0.57 | 0.72 |
| Paclitaxel + Ganetespib | HR-HER2- | 0.81 | 0.78 |
| Paclitaxel + Ganitumab | HR-HER2- | 0.85 | 0.85 |
| Paclitaxel + Ganitumab | HR+HER2- | 0.55 | 1 |
| Paclitaxel + MK-2206 | HR+HER2- | 0.7 | 1 |
| Paclitaxel + MK-2206 | HR-HER2- | 0.33 | 1 |
| Paclitaxel + Neratinib | HR-HER2- | 0.94 | 0.77 |
| Paclitaxel + Neratinib | HR+HER2- | 1 | 0.71 |
| Paclitaxel + Neratinib | HR+HER2+ | 0.89 | 1 |
| Paclitaxel + Pembrolizumab | HR+HER2- | 0.68 | 0.8 |
| Paclitaxel + Pertuzumab + Trastuzumab | HR+HER2+ | 0.71 | 1 |
| Paclitaxel + Trastuzumab | HR+HER2+ | 1 | 1 |
| T-DM1 + Pertuzumab | HR+HER2+ | 1 | 1 |
